# Fast, flexible gene cluster family delineation with IGUA

**DOI:** 10.1101/2025.05.15.654203

**Authors:** Martin Larralde, Josefin Blom, Hadrien Gourlé, Laura M. Carroll, Georg Zeller

**Author notes:** Corresponding authors: Laura M. Carroll,; Georg Zeller. Joint Authors (Martin Larralde, Josefin Blom, Laura M. Carroll, Georg Zeller).

## Abstract

Prokaryotic genomes harbor a variety of functional elements encoded as contiguous multi-gene clusters, with biosynthetic gene clusters (BGCs, genetic determinants of secondary metabolite biosynthesis) serving as a notable example. In a typical workflow, BGCs are clustered into Gene Cluster Families (GCFs), units that group BGCs encoding similar biosynthetic pathways together. However, existing methods cannot readily scale to massive datasets and cannot be used for GCF delineation tasks beyond BGC clustering. Here, we present IGUA (Iterative Gene clUster Analysis; https://github.com/zellerlab/IGUA), a scalable, flexible GCF delineation method for genomic segments with multi-gene architectures. On a BGC clustering task, IGUA is ≥10x faster than the state-of-the-art (BiG-SCAPE/BiG-SLiCE), without sacrificing accuracy. To highlight its scalability, we use IGUA to cluster >2.8 million BGCs from ≈1 million prokaryotic genomes in <18 hours (*n* = 2,829,071 BGCs to 56,960 GCFs). To showcase its utility beyond BGC clustering, we use IGUA to cluster (i) secretion systems and (ii) prophages into GCFs (*n* = 10,576 and 356,776 gene clusters to 2,744 and 213,699 GCFs, respectively). Overall, IGUA represents a versatile GCF delineation tool with unmatched computational efficiency and flexibility, enabling (meta)genomic mining applications at unprecedented scales.

**GRAPHICAL ABSTRACT:** 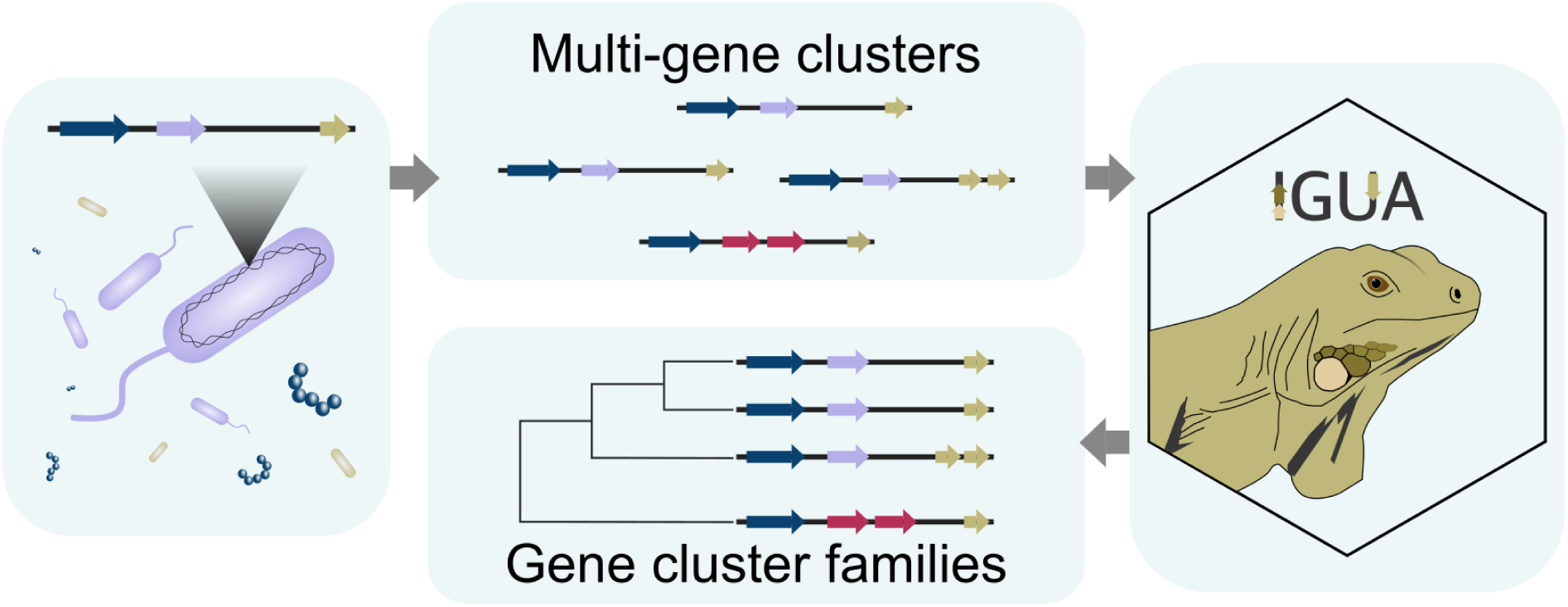

## INTRODUCTION

In prokaryotic genomes, genome organization is indicative of function, and proteins that can be functionally grouped into the same pathway tend to be encoded as genes that are in close proximity to each other in the genome (1, 2). A variety of bioinformatic methods/tools have been developed for detecting multi-gene clusters in prokaryotic (meta)genomes (3–5), including specialist tools designed to detect specific types of prokaryotic gene clusters, such as genomic islands (6–9), secretion systems (10), prophage (11–16), phage defense systems (17–19), and biosynthetic gene clusters (BGCs) (20–24). Regardless of their function, in order to study the evolution and diversity of these prokaryotic gene clusters at scale, bioinformatic methods that account for their multi-gene architectures are paramount.

BGCs (biosynthetic gene clusters) are one specific class of prokaryotic gene cluster and an increasingly sought-after target of (meta)genomic mining efforts (25–28). BGCs equip their prokaryotic host with the ability to produce secondary metabolites (25, 28). Also known as specialized metabolites or natural products, secondary metabolites are small, auxiliary molecules produced by microbes for fitness advantages within their ecological niche (28). Beyond their ecological functions, microbial secondary metabolites and their derivatives have important roles in medicine and industry, particularly within the realm of drug discovery, where they have found success as critically important antibacterial, antifungal, immunosuppressive, and chemotherapeutic agents (25, 26, 29, 30).

While a range of data modalities can be leveraged for novel secondary metabolite discovery (e.g., metabolomic, proteomic, and/or phenotypic data), BGC mining methods have been particularly promising; they are relatively fast, inexpensive, and can utilize the exponentially growing amount of (meta)genomic data available in public databases (25–27). In a typical novel BGC discovery workflow, BGCs are first detected in a set of microbial genomes (e.g., using antiSMASH, arguably the most popular BGC detection tool for over a decade) (20). BGCs can then be clustered into Gene Cluster Families (GCFs), units that attempt to group BGCs encoding similar biosynthetic pathways together (27). GCFs can then be used to e.g. identify and filter out redundant BGCs, assign functions to unknown BGCs, and identify putative novel BGCs (27).

Several GCF delineation tools have been developed, with BiG-SCAPE (27) and BiG-SLiCE (31) being arguably the most established. BiG-SCAPE works by taking BGC nucleotide sequences (from antiSMASH and/or MIBiG, a database of experimentally validated BGCs) (20, 27, 28) as input, translating them into protein sequences, and representing each as a string of Pfam protein domains (27, 32). BGCs are then clustered into GCFs using a combination of three distance metrics (Jaccard index, adjacency index, and domain sequence similarity index), weighted differently depending on the biosynthetic class of the BGCs (27). While accurate and highly sensitive,

BiG-SCAPE is slow, and its quadratic runtime complexity prevents it from scaling to large datasets (i.e., >10^4^ BGCs) (31). To overcome this limitation, BiG-SLiCE was designed to delineate GCFs in a similar fashion, but with algorithmic changes that sacrifice some accuracy in favor of speed (31).

Despite achieving near-linear time complexity (31), BiG-SLiCE is still too slow for many large-scale applications. For example, clustering >1.2 million BGCs into GCFs on a 36-core CPU server took BiG-SLiCE nearly 10 days (31). Instead of flat text files, BiG-SLiCE outputs results in the form of interactive HTML reports (31), making it challenging to use results for downstream tasks outside of the BiG-SLiCE ecosystem. Further, both BiG-SCAPE and BiG-SLiCE are dependent on the presence of Pfam protein domains for clustering (27, 31, 32); novel gene clusters with low-to-no Pfam domain coverage thus cannot be queried within this framework. Finally, both

BiG-SCAPE and BiG-SLiCE were created to work optimally with the output of antiSMASH and/or MIBiG (20, 27, 28, 31). The increasing availability of alternative BGC detection tools (e.g., machine learning [ML]-based methods) (21, 22, 24) necessitates the development of more flexible GCF delineation tools. Overall, as BGC mining efforts increase in size and scope, it is apparent that existing GCF delineation methods are insufficient in terms of scalability, computational efficiency, and flexibility.

Here, we present Iterative Gene clUster Analysis (IGUA; https://github.com/zellerlab/IGUA), a scalable, flexible GCF delineation method for genomic segments with multi-gene architectures. Created with large-scale users in mind, IGUA outperforms gold-standard GCF delineation tools in terms of time and memory requirements, without sacrificing accuracy. On a BGC clustering task, we show that IGUA is >47 and >9x faster than BiG-SCAPE and BiG-SLiCE, respectively, and to showcase its scalability, we use IGUA to cluster >2.8 million BGCs from nearly 1 million prokaryotic genomes in under 18 hours. Finally, we show that IGUA can be used for clustering tasks beyond the BGC space by clustering (i) secretion system and (ii) prophage gene clusters into GCFs. Taken together, IGUA represents a versatile GCF delineation tool with unprecedented scalability and flexibility, which can be leveraged in large-scale, data-driven (meta)genomic mining applications.

## MATERIALS AND METHODS

### Implementation of IGUA

IGUA was designed to cluster gene clusters (e.g., BGCs) into GCFs using three major steps: (i) a fragment removal step, followed by (ii) nucleotide-level deduplication and (iii) protein-level clustering steps (Figure 1). Briefly, (i) the “mmseqs linclust” command in MMseqs2 (at the time of publication, release 15-6f452) (33) identifies and removes fragmented gene clusters, which are substrings of a larger gene cluster, using the following parameters (i.e., low-coverage/high-identity mode; Figure 1A): *E*-value = 0.001, sequence identity = 0.85, coverage = 1, cluster mode = 0, coverage mode = 1, spaced *k*-mer mode = 0. After removing gene cluster fragments, (ii) “mmseqs linclust” is used a second time to deduplicate closely related gene clusters at the nucleotide (DNA) level, using the following parameters (i.e., high-coverage/medium-identity mode; Figure 1A): *E*-value = 0.001, sequence identity = 0.6, coverage = 0.5, cluster mode = 0, coverage mode = 0, spaced *k*-mer mode = 0.

**Figure 1.**
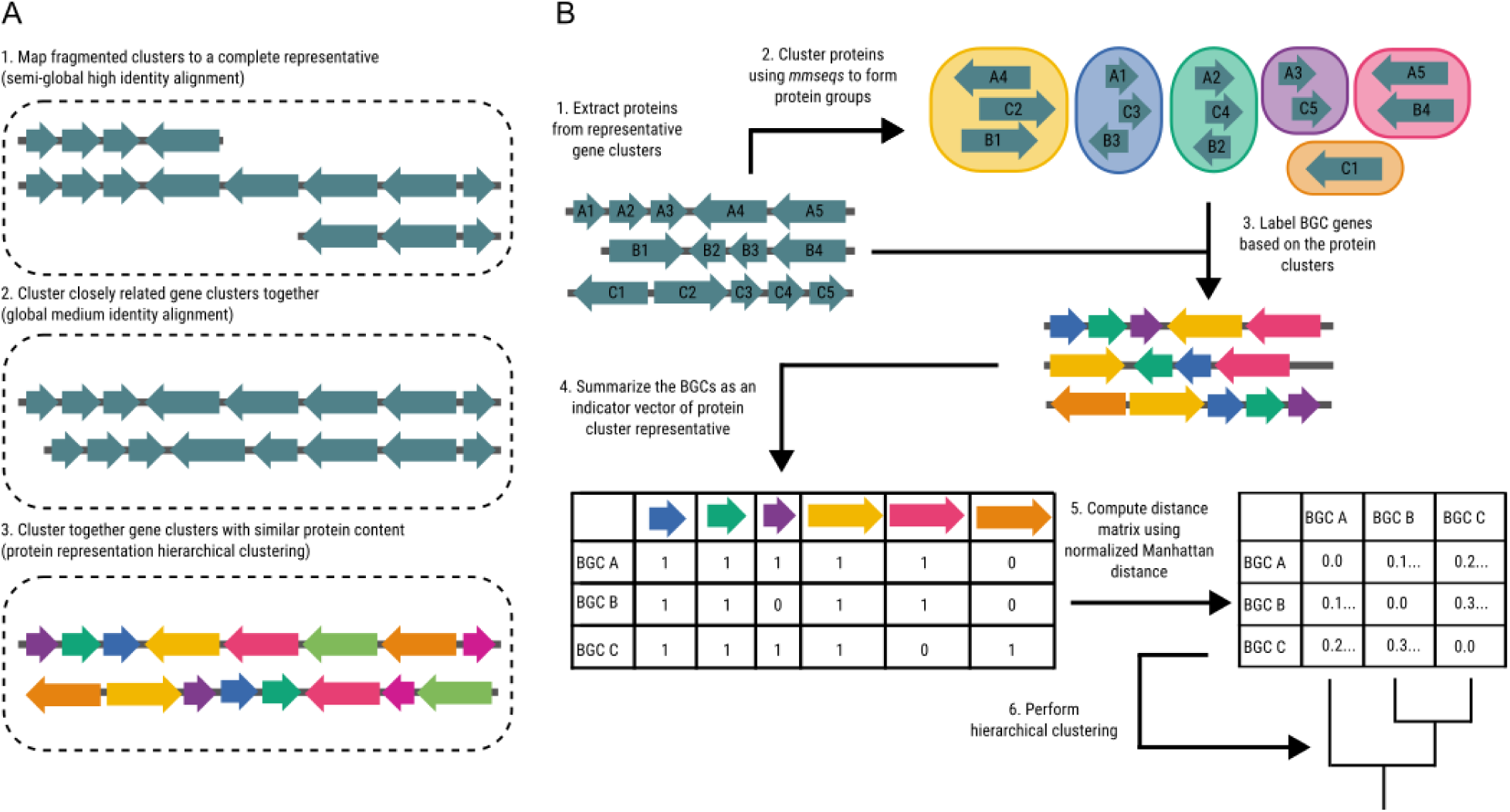
Overview of IGUA’s iterative Gene Cluster Family (GCF) delineation pipeline. **(A)** IGUA clusters gene clusters (e.g., biosynthetic gene clusters [BGCs]) into GCFs using three major steps: (1) a fragment removal step, (2) a nucleotide (DNA)-level deduplication step, and (3) a protein (amino acid)-level clustering step. **(B)** IGUA’s protein-level clustering step (step 3) in detail.

For (iii) the protein clustering step, protein (amino acid) sequences of MMseqs2 representative gene clusters from step (ii) are clustered using “mmseqs linclust” (parameters: *E*-value = 0.001, coverage = 0.9, coverage mode = 1, sequence identity = 0.5), resulting in a set of “protein groups” (Figure 1A). Protein group counts are then used to populate a sparse protein composition matrix, in which each row represents a gene cluster, each column represents a protein group, and each cell contains the number of proteins in each gene cluster assigned to each protein group (Figure 1B). The length of each protein group representative is recorded (to be used as weights in the subsequent pairwise distance computation). Weighted Manhattan distances are then calculated between all gene clusters using a fast parallel algorithm designed for compressed-sparse-row (CSR) matrices (Figure 1B). The distances between two clusters are normalized using the sum of their weights. Gene clusters are then partitioned into GCFs using hierarchical clustering, using the linkage function from the kodama library (at the time of publication, v0.2.0; https://crates.io/crates/kodama) and a user-provided agglomerative clustering method. Together, this results in a set of GCFs, as well as a set of representative gene clusters (one per GCF; i.e., the longest gene cluster from step ii within the GCF; Figure 1B).

### Optimization of IGUA parameters on a manually curated dataset

To provide users with optimal default settings for IGUA’s two user-specified parameters (i.e., hierarchical clustering linkage method and clustering distance), we used IGUA to cluster BGCs into GCFs, testing a range of parameter combinations on a set of manually curated BGC-to-GCF mappings. Briefly, a manually curated dataset consisting of 313 experimentally validated BGCs associated with 376 chemical compounds (i.e., the same dataset used to benchmark BiG-SCAPE) was extracted from MIBiG v1.3 (downloaded from BiG-SCAPE’s GitHub repository; https://github.com/medema-group/BiG-SCAPE/tree/master/Annotated_MIBiG_reference) (27, 28). To mimic the analysis performed in the original BiG-SCAPE paper (27), two of the classes in the manually curated dataset (microcins and lanthipeptides I) were omitted prior to clustering, resulting in a data set consisting of 304 experimentally validated BGCs associated with 366 chemical compounds. Additionally, BGCs with >1 GenBank record per file were removed post-clustering (as IGUA would recognize them as different BGCs, while BiG-SCAPE would not), resulting in a final data set of 292 BGCs associated with 352 chemical compounds (referred to hereafter as the “manually curated dataset”; Supplementary Table S1).

To select optimal default clustering parameters for IGUA, BGCs in the manually curated dataset were clustered into GCFs using IGUA v0.1.0, testing all combinations of the following parameters: (i) linkage method, set to one of “average”, “centroid”, “ward”, “weighted”, “complete”, “median”, or “single”; (ii) clustering distance, set from 0.01 to 1.0 in 0.01 intervals (700 total runs representing 700 total parameter combinations). For each IGUA run, singleton GCFs were removed (as was done in the original BiG-SCAPE study, so as to not skew subsequent calculations of GCF purity) (27). GCFs produced via IGUA were then compared to the manually curated GCFs (the latter representing “ground truth” GCFs) in R v4.1.2 (https://www.R-project.org/, accessed 23 December 2024), and the following metrics were calculated: (i) adjusted Rand index (ARI), (ii) adjusted mutual information (AMI), and (iii) normalized mutual information (NMI) scores (aricode package v1.0.3; https://CRAN.R-project.org/package=aricode, accessed 23 December 2024); (iv) purity (funtimes package v9.1; https://CRAN.R-project.org/package=funtimes, accessed 23 December 2024); (v) total number of GCFs. A combination of linkage method = “average” and clustering distance = 0.80 was determined to yield optimal results (Figure 2A, Supplementary Figure S1); these parameters were set as the default parameters in IGUA v0.1.0 and were used in all subsequent steps unless otherwise stated.

**Figure 2.**
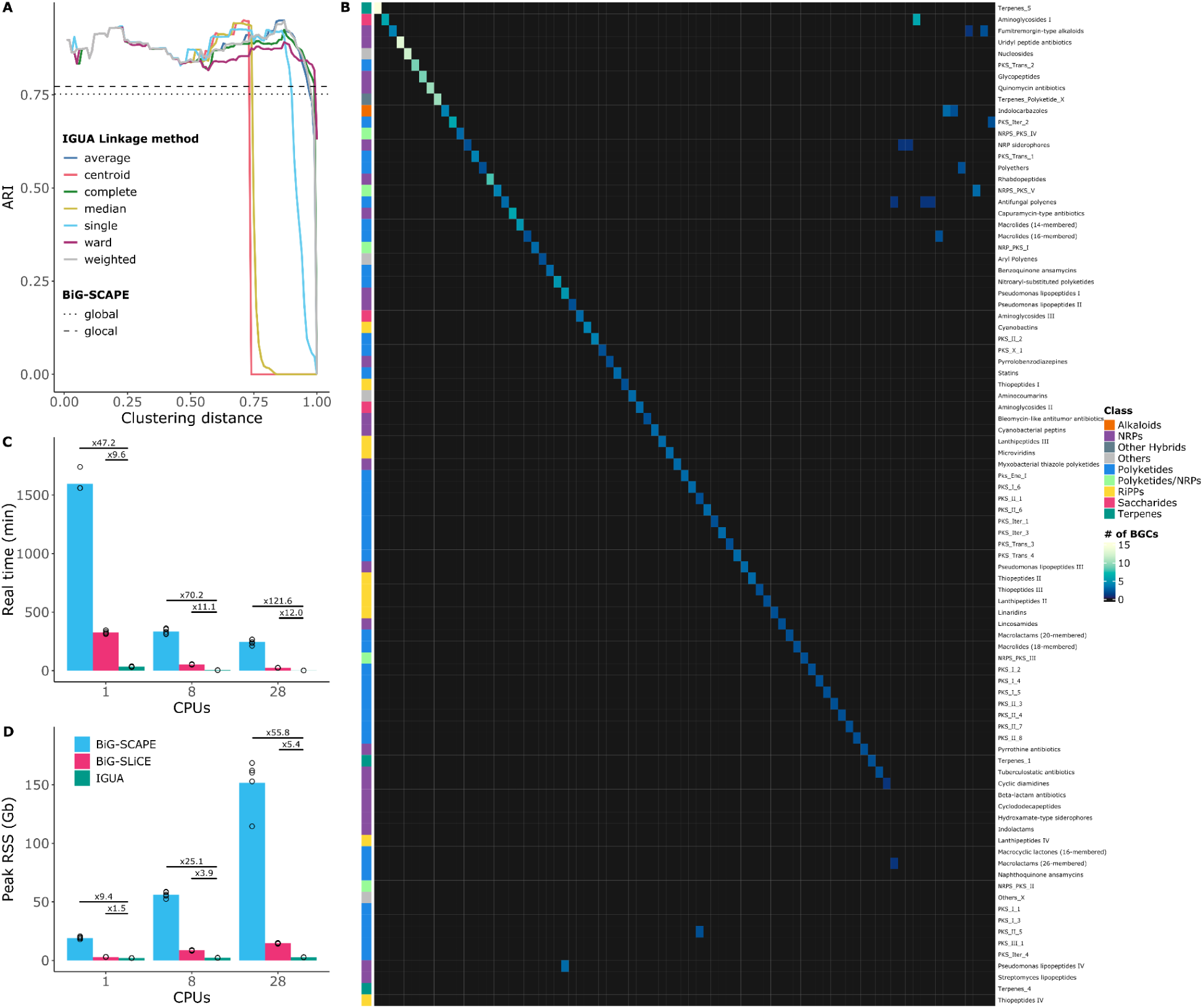
IGUA is ≥10x faster than the state-of-the-art, without sacrificing accuracy. **(A)** Adjusted Rand index (ARI; Y-axis) of IGUA GCFs relative to manually curated, “ground truth” GCFs for each tested parameter combination (clustering distance, X-axis; hierarchical clustering linkage method, line color). Horizontal dashed lines denote ARI values achieved by BiG-SCAPE in “global” or “glocal” mode (where a theoretical maximum of 1.0 = the optimal ARI value, corresponding to GCFs that are identical to the ground truth). A clustering distance of 0.80 and the “average” linkage method were selected as IGUA’s default parameters. For additional clustering metrics, see Supplementary Figure S1. **(B)** Heatmap comparing manually curated, “ground truth” GCFs (rows) to GCFs delineated using IGUA (linkage method = “average”, clustering distance = 0.80; columns). Cells are shaded according to the number of overlapping BGCs within a GCF. The color strip to the left of the heatmap denotes the biosynthetic class of each manually curated GCF. NRPs, non-ribosomal peptide synthetases; RiPPs, ribosomally synthesized and post-translationally modified peptides. **(C)** Average time (real/“wall clock” time, in minutes; Y-axis) and **(D)** memory usage (peak resident set size [RSS] in Gb; Y-axis) for BiG-SCAPE (blue), BiG-SLiCE (pink), and IGUA (green), using 1, 8, or 28 CPUs (X-axes). For each tool, GCF delineation was carried out using antiSMASH BGCs from five diverse subsets of the proGenomes3 database (≈5,000 genomes each). Dots overlapping each bar denote raw values for each proGenomes3 subset.

### Evaluation of IGUA and BiG-SCAPE accuracy on a manually curated dataset

To evaluate the GCF delineation accuracy of IGUA relative to the state-of-the-art, (i) BiG-SCAPE v1.1.5 (27) and (ii) IGUA v0.1.0 were used to cluster BGCs from the manually curated dataset into GCFs (Supplementary Table S1, see section “Optimization of IGUA default parameters on a manually curated dataset” above; note that BiG-SLiCE was not included here, as BiG-SCAPE has already been determined to be more accurate than BiG-SLiCE) (31). For (i) BiG-SCAPE v1.1.5, BGCs were clustered into GCFs in two separate runs, using the following parameters (i.e., the same parameters used in the original BiG-SCAPE paper): hybrid mode disabled (--hybrids-off; both runs), a cutoff value of 0.75 (--cutoffs 0.75; both runs), and either glocal or global alignment mode (--mode glocal or --mode global; run 1 and run 2, respectively; Supplementary Tables S2-S3) (27). For (ii) IGUA v0.1.0, BGCs were clustered into GCFs using optimal parameters (linkage method = “average” and clustering distance = 0.80, see section “Optimization of IGUA default parameters on a manually curated dataset” above; Supplementary Table S4). Post-clustering, singleton GCFs (<2 BGCs) were excluded from further analysis (as was done in the original BiG-SCAPE paper) (27). BiG-SCAPE and IGUA GCFs were each compared to the manually curated GCFs (representing “ground truth” GCFs) as described above (see section “Optimization of IGUA parameters on a manually curated dataset”; Figure 2B, Supplementary Figure S2, Supplementary Table S1).

### Comparison of IGUA and BiG-SCAPE GCFs on a *Streptomyces* dataset

In addition to the manually curated dataset (see section “Evaluation of IGUA and BiG-SCAPE accuracy on a manually curated dataset” above), a dataset consisting of complete *Streptomyces* genomes was used to compare GCFs delineated by IGUA to those delineated by BiG-SCAPE (similar to the *Streptomyces* dataset used in the original BiG-SCAPE paper; note that the exact *Streptomyces* dataset described in the BiG-SCAPE paper could not be used, as not all genomes/accession numbers were provided) (27). Briefly, GenBank assemblies for all complete *Streptomyces* genomes in NCBI’s Assembly database (34) were downloaded (*n* = 492 complete genomes, accessed 9 December 2023). antiSMASH v6.1.1 (35) was used to detect BGCs in each *Streptomyces* genome, using Prodigal in metagenomic mode as the genefinding tool (--genefinding-tool prodigal-m) and two CPUs (-c 2; remaining parameters were set to their defaults) (36). The resulting antiSMASH BGCs (*n* = 15,298) were clustered into GCFs using: (i) BiG-SCAPE v1.1.5, with “glocal” mode (--mode glocal), singletons included (--include_singletons), hybrid mode disabled (--hybrids_off), and MIBiG v3.1 included (--mibig) (37); (ii) IGUA v0.1.0, which was run with optimal parameters (linkage method = “average” and clustering distance = 0.80) and MIBiG v3.1 BGCs included (*n* = 17,800 BGCs total, corresponding to 2,502 and 15,298 BGCs from MIBiG v3.1 and antiSMASH, respectively). AMI, NMI, ARI, and purity were calculated relative to BiG-SCAPE clustering results as described above (see section “Optimization of IGUA parameters on a manually curated dataset”; Supplementary Figure S3, Supplementary Tables S5-S6).

To further assess IGUA’s speed relative to input dataset size, the full set of *Streptomyces* antiSMASH BGCs (with MIBiG v3.1 BGCs included) was down-sampled to (i) 1,000, (ii) 2,500, (iii) 5,000, and (iv) 10,000 BGCs in triplicate (*n* = 12 total down-sampled *Streptomyces* datasets) using the random.sample() function from the random module of Python v3.10.4. Each down-sampled *Streptomyces* dataset was clustered using IGUA v0.1.0 (8 CPUs and optimal parameters, i.e., linkage method = “average” and clustering distance = 0.80), with runtime logged using Nextflow v24.04.2 (i.e., by enabling the “report” option in Nextflow’s configuration file; Supplementary Figure S4, Supplementary Tables S7-S8) (38).

### Speed and memory benchmarking using species representative genomes

To evaluate speed and memory requirements for each of BiG-SCAPE v1.1.5 (27), BiG-SLiCE v1.1.0 (31), and IGUA v0.1.0, BGCs from five diverse subsets of the proGenomes3 database (39) were used. Briefly, antiSMASH v6.1.1 was used to detect BGCs in all proGenomes3 representative genomes as described above (*n* = 41,777 genomes; see section “Comparison of IGUA and BiG-SCAPE GCFs on a *Streptomyces* dataset”). Genome Taxonomy Database (GTDB) taxonomic assignments (40) for all proGenomes3 representative genomes were downloaded from proGenomes3 (https://progenomes.embl.de/data/proGenomes3_specI_lineageGTDB.tab.bz2; accessed 17 October 2022) (40, 41). To ensure that each subset of the proGenomes3 representative genomes contained diverse, non-redundant genomes, one specI cluster (i.e., species) (41) per GTDB genus was sampled, and from this, 5,000 specI clusters were randomly selected. For each randomly selected specI cluster, the specI representative genome was identified (per proGenomes3) and selected. This process was repeated five times to generate five samples of approximately 5,000 genomes each (some specI clusters lacked representative genomes; in these rare cases, the specI cluster was not included in the subset). Overall, this resulted in five subsets, each containing antiSMASH BGCs from ≈5,000 diverse genomes, each representing a distinct prokaryotic genus (Supplementary Table S9).

For each of the five subsets, all antiSMASH BGCs from the sampled representative genomes were clustered with (i) BiG-SCAPE v1.1.5, (ii) BiG-SLiCE v1.1.0, and (iii) IGUA v0.1.0. For (i) BiG-SCAPE, antiSMASH BGCs were clustered alongside BGCs from the MIBiG database (--mibig; v3.1), using glocal mode (--mode glocal), the “include singletons” option enabled (--include_singletons), and “hybrid mode” disabled (--hybrids-off). For (ii) BiG-SLiCE, default settings were used to cluster all antiSMASH BGCs into GCFs (MIBiG BGCs were not included, as BiG-SLiCE did not accept MIBiG-formatted BGCs). For (iii) IGUA, antiSMASH BGCs detected in each subset of genomes were concatenated with BGCs from MIBiG v3.1; IGUA was then used to cluster each concatenated subset using optimal settings (linkage method = “average” and clustering distance = 0.80). For each of the five subsets, all three methods were used to cluster BGCs into GCFs using 1, 8, or 28 CPUs. For each run, runtime and memory requirements were logged by enabling the “report” option in the Nextflow configuration file (Figure 2CD; Supplementary Table S10).

### Clustering of >2.8 million GECCO BGCs from >900,000 prokaryotic genomes

To showcase its speed and scalability, IGUA was used to cluster all BGCs detected in the proGenomes3 database (39) into GCFs. Briefly, GenBank records for all proGenomes3 BGCs were downloaded (*n* = 2,826,569 total BGCs from 907,388 total genomes, detected using GECCO, a ML-based BGC detection method, as described previously; https://progenomes.embl.de/data/progenomes3_gecco_clusters.gbk.gz) (21, 39). IGUA v0.1.0 was used to cluster all 2,826,569 proGenomes3 GECCO BGCs, plus 2,502 experimentally validated BGCs from MIBiG v3.1, into GCFs (72 CPUs, linkage method = “average”, and clustering distance = 0.80; Supplementary Table S11).

Runtime and memory usage were logged by enabling the “report” option in the Nextflow configuration file.

To visualize proGenomes3 GCFs in two-dimensional space, a UMAP (42) was constructed using GCF representative BGCs output by IGUA (*n* = 56,960 GCF representative BGCs). To construct the UMAP, pyHMMER v0.8.2 (43) was used to annotate each IGUA GCF representative BGC (28 CPUs and default settings), using Pfam v35.0 profile hidden Markov models (pHMMs; downloaded from the Pfam FTP site (https://ftp.ebi.ac.uk/pub/databases/Pfam/releases/Pfam35.0/Pfam-A.hmm.gz) (32).

From this, a protein domain composition matrix was computed (via https://github.com/c20josbl/igua_manuscript_documentation/blob/main/umap_bgcs.py) and extracted for UMAP construction. The UMAP was produced in R v4.3.1, using the packages Seurat v5.0.2 (44), Signac v1.12.0 (45), reticulate v1.35.0 (https://CRAN.R-project.org/package=reticulate, accessed 23 December 2024), and clustree v0.5.1 (46). Briefly, the compositions matrix was imported using the scipy.sparse$load_npz function and made into a Seurat object (CreateSeuratObject(), assay = “peaks”) via intermediate conversions to an RDS object, followed by a chromatin assay (CreateChromatinAssay()). This was followed by normalization and selection of top features (RunTFIDF() and FindTopFeatures(), respectively, with min.cutoff =”q0” for the latter) prior to dimensional reduction with singular value decomposition (SVD) using RunSVD(). Thereafter, RunUMAP() was run on the top 30 SVD dimensions, with reduction = “lsi”, n.neighbors = 40L, and min.dist = 0.2.

FindNeighbours() was used to create a nearest-neighbour graph (reduction = “lsi”, dims = 1:30). For selection of an appropriate clustering resolution, a clustree was created for resolutions 0.10-0.90 with 0.05 increments (Supplementary Figure S5). Finally, the clusters of the UMAP were created using FindClusters() (resolution = 0.65, algorithm = 3). The resulting Seurat object was saved as pfams_proG3.RDS using saveRDS(). The predicted biosynthetic class(es) assigned to each BGC (obtained using GECCO as described previously) (39) were extracted from each BGC’s GenBank record. When >1 protocluster was present, the BGC was labelled as “Mixed”. The predicted biosynthetic class(es) assigned to the majority of BGCs in each GCF was attached to the pfams_proG3.RDS object using AddMetaData() (Supplementary Figure S5, Supplementary Table S12).

### Clustering of secretion system and prophage gene clusters

To showcase its utility for tasks beyond BGC clustering, we used IGUA to cluster (i) secretion system and (ii) prophage gene clusters into GCFs. For (i) secretion system clustering, metadata for all secretion systems in TXSSdb (10) (i.e., nucleotide accession numbers, start/end coordinates, and secretion system type classifications) were downloaded (*n* = 11,939 total secretion system entries; http://macsydb.web.pasteur.fr/txssdb/_design/txssdb/index.html, accessed 27 September 2024). Secretion systems classified by TXSSdb as “single_locus” secretion systems were used in subsequent steps (*n* = 10,784 TXSSdb single-locus secretion systems). The “entrez_fetch” command in the rentrez package (v1.2.3) (47) in R v4.3.2 was used to query NCBI’s Nucleotide database (48) for each single-locus secretion system, using NCBI Nucleotide accession numbers and secretion system start/end coordinates provided by TXSSdb. This resulted in a total of 10,576 valid GenBank files (one per single-locus secretion system), which were concatenated into a single GenBank file using BioPython v1.84 (49). IGUA v0.1.0 was used to cluster all 10,576 TXSSdb single-locus secretion systems into GCFs (linkage method = “average”, clustering distance = 0.80; Supplementary Table S13). The “entrez_search” and “entrez_summary” commands in rentrez were further used to download the host bacterium genome for each secretion system, and each host bacterium genome underwent taxonomic classification using GTDB release214/GTDB-Tk v2.3.2 (i.e., via GTDB-Tk’s “classify_wf” workflow with --cpus 28 and default settings; Supplementary Table S14) (40, 50).

A UMAP of the IGUA GCF representative secretion systems was constructed as described above (see section “Clustering of >2.8 million GECCO BGCs from >900,000 prokaryotic genomes”), with the following alterations: the protein domain composition matrix was computed using https://github.com/c20josbl/igua_manuscript_documentation/blob/main/umap_secretion.py, the RunUMAP() options min.dist and n.neighbours were not changed from default settings, and the clustering resolution for FindClusters() was set to 0.35. The Seurat object was saved as pfams_secretion.RDS, and the attached metadata consisted of (a) secretion system types (provided by TXSSdb), and (b) host bacterium taxonomic classifications (per GTDB-Tk; Supplementary Figure S6, Supplementary Tables S14-S15). This process was repeated a second time, but with the clustering resolution for FindClusters() set to 0.80 (the default; Supplementary Figure S7).

For (ii) prophage clustering, all prophage sequences in Prophage-DB (51) were downloaded (i.e., bacterial_host_prophages.tar.gz, archaeal_host_prophages.tar.gz, unknown_host_prophages.tar.gz, and all corresponding metadata; https://doi.org/10.5061/dryad.3n5tb2rs5, accessed 26 November 2024). Prokka v1.14.6 (52) was used to annotate each prophage using its respective FASTA file as input, the “Viruses” database (--kingdom Viruses), and the “--addgenes” and “--compliant” options enabled (remaining options were set to their defaults). Annotated GenBank files produced by Prokka (*n* = 356,776 files, one per Prophage-DB prophage) were concatenated into a single GenBank file and clustered into GCFs using IGUA (linkage method = “average”, clustering distance = 0.80; Supplementary Table S16). Due to the large number of resulting GCFs (*n* = 213,699), IGUA was run a second time, using average linkage and a higher clustering distance threshold (i.e., 0.99, resulting in 106,749 GCFs; Supplementary Table S17). IGUA GCF representatives identified using this higher threshold (0.99) were visualized via a UMAP, which was constructed as described above, with the following changes: the protein domain composition matrix was computed using https://github.com/c20josbl/igua_manuscript_documentation/blob/main/umap_prophage.py, and the options min.dist and n.neighbours for RunUMAP() and clustering resolution for FindClusters() were not altered from default settings. The SeuratObject was saved as pfams_prophage.RDS, and the attached metadata consisted of host bacterium taxonomic classifications (provided by Prophage-DB; Supplementary Figure S8, Supplementary Table S18).

### Data visualization

Plots were constructed in R v4.3.1, using the following packages (all URLs accessed 23 December 2024): readxl v1.4.3 (https://CRAN.R-project.org/package=readxl), tidyverse v2.0.0 (53), khroma v1.13.0 (https://packages.tesselle.org/khroma/), ggnewscale v0.4.10 (https://CRAN.R-project.org/package=ggnewscale), ggpubr v0.6.0 (https://CRAN.R-project.org/package=ggpubr), ComplexHeatmap v2.18.0 (54), ggplotify v0.1.2 (https://CRAN.R-project.org/package=ggplotify), ggalluvial v0.12.5 (http://corybrunson.github.io/ggalluvial/), ggmagnify v0.4.1.9000 (https://github.com/hughjonesd/ggmagnify), ggsignif v0.6.4 (55), Signac v1.12.0 (45), Seurat v5.0.2 (44), clustree v0.5.1 (46), and png v0.1-8 (https://CRAN.R-project.org/package=png).

## RESULTS

### IGUA: accurate, scalable GCF delineation via iterative clustering

The IGUA pipeline is composed of three major steps, which are designed to tackle three major problems associated with GCF delineation (Figure 1). The first step, (i) a fragment removal step, uses MMseqs2 in low-coverage/high-identity mode to remove sequences that are fragments (subsequences or substrings) of a larger gene cluster (Figure 1A).

This reduces computation time/memory requirements needed in later steps, particularly for metagenomic datasets, which are often composed of highly redundant, fragmented metagenome-assembled genomes (MAGs).

The second step, (ii) nucleotide-level deduplication, uses MMseqs2 in high-coverage/medium-identity mode to group highly similar (e.g., nearly identical) gene clusters together, producing a set of representative gene clusters (Figure 1A). This further reduces computation time/memory requirements needed in the third step, (iii) protein-level clustering. In this final step, all protein sequences undergo unsupervised clustering via MMseqs2 to produce protein groups, which are then used to produce a sparse matrix, in which each gene cluster is encoded as a vector of protein group representatives (Figure 1B). Pairwise distances between gene clusters are computed using weighted and normalized Manhattan distances, in order to account for variability in protein length and total cluster length between each cluster pair (Figure 1B). In other words, the distance computation between gene clusters is weighted using the length of each protein group representative; this ensures that the absence of a larger gene (e.g., a core biosynthetic gene in a non-ribosomal peptide synthetase) is weighted more heavily than the absence of a smaller gene (e.g., a regulatory gene). Finally, hierarchical clustering is then used to delineate gene clusters into GCFs (Figure 1). Altogether, IGUA’s iterative clustering approach removes redundancy early on (i.e., in the fragment removal and nucleotide-level deduplication steps) to reduce time and memory usage later (i.e., in the more computationally intensive, protein-level clustering step; Figure 1).

### IGUA outperforms the state-of-the-art in terms of compute time and memory usage, without sacrificing accuracy

IGUA uses two user-specified parameters for GCF delineation: (i) linkage method (for hierarchical clustering/dendrogram construction), and (ii) clustering distance (the distance at which the dendrogram is “cut” to produce GCFs; Figure 1B). To provide users with optimal default parameters, we used IGUA to cluster BGCs into GCFs, testing a range of parameter combinations on a set of manually curated, “ground-truth” GCFs (i.e., the dataset used to benchmark and validate BiG-SCAPE) (27). After considering GCF composition/purity and the total number of GCFs produced, we selected (i) average linkage hierarchical clustering with (ii) a clustering distance threshold of 0.80 as the optimal default parameter combination for IGUA (Figure 2A, Supplementary Figure S1; see section “Optimization of IGUA parameters on a manually curated dataset” above). Notably, using these selected parameters, IGUA performed slightly better than BiG-SCAPE on this manually curated, “ground truth” BiG-SCAPE validation dataset (Figure 2B, Supplementary Figure S2, Supplementary Tables S2-S4). On the *Streptomyces* dataset (i.e., a set of 15,298 antiSMASH BGCs from 492 complete *Streptomyces* genomes, similar to the *Streptomyces* dataset used in the BiG-SCAPE validation/benchmarking paper), both IGUA and BiG-SCAPE produced similar GCFs (ARI = 0.83 and 0.82 with singleton GCFs included and removed, respectively; Supplementary Figure S3, Supplementary Tables S5-S6, see section “Comparison of IGUA and BiG-SCAPE GCFs on a *Streptomyces* dataset” above). Overall, IGUA’s GCF delineation accuracy is on par with the state-of-the-art for BGC clustering tasks.

Beyond validating the accuracy of IGUA’s GCF delineation method, we further benchmarked IGUA’s speed and memory usage relative to BiG-SCAPE (the state-of-the-art method in terms of accuracy) and BiG-SLiCE (the state-of-the-art method in terms of speed; Figure 2CD). To accomplish this, each of the three methods (BiG-SCAPE, BiG-SLiCE, and IGUA) was used to cluster five sets of BGCs from ≈5,000 diverse prokaryotic genomes (each representing a unique genus) into GCFs, using 1, 8, or 28 CPUs (Figure 2CD, Supplementary Table S10). Using 1 CPU, IGUA was 47 and 10x faster on average than BiG-SCAPE and BiG-SLiCE, respectively; as the number of CPUs increased, these differences in runtime increased (Figure 2C). Similar results were observed for memory usage (Figure 2D, Supplementary Table S10). Taken together, IGUA represents an accurate GCF delineation method, which is over an order of magnitude faster than the state-of-the-art and substantially more memory efficient.

### IGUA readily scales to millions of biosynthetic gene clusters

To showcase the speed, scalability, and flexibility of IGUA, we delineated GCFs among >2.8 million BGCs, detected in nearly 1 million prokaryotic genomes using GECCO, a ML-based BGC detection method (21, 39). Using 72 CPUs, IGUA clustered 2,829,071 BGCs (2,826,569 GECCO BGCs detected in 907,388 genomes from the proGenomes3 database, plus 2,502 experimentally validated BGCs from the MIBiG database) (39) into 56,960 GCFs in 17 hours and 22 minutes (Figure 3A). Notably, an overwhelming proportion of the resulting GCFs did not contain any experimentally validated MIBiG BGCs (55,071 of 56,960 total GCFs, 96.7%), indicating that there are massive portions of the prokaryotic biosynthetic landscape, which have never been queried experimentally (Figure 3A, Supplementary Figure S5, Supplementary Tables S11-S12). Overall, IGUA can readily scale to massive datasets consisting of millions of BGCs, including BGCs detected using newer ML methods.

**Figure 3.**
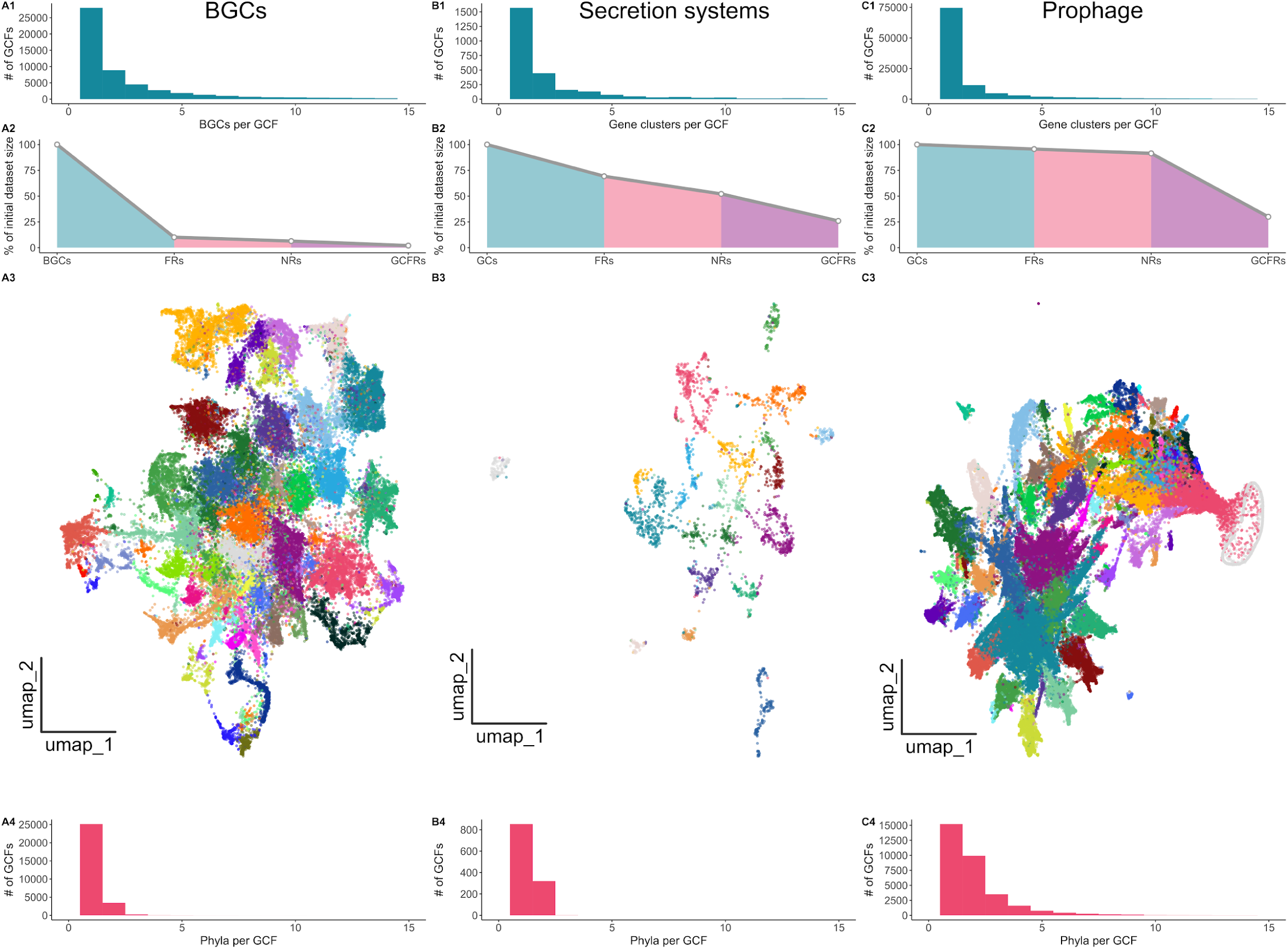
IGUA is a flexible GCF delineation tool, which can be used for tasks beyond BGC clustering. IGUA was used to cluster **(A)** BGCs, **(B)** secretion system, and **(C)** prophage gene clusters into GCFs, with plots denoting the following: (**1**) histograms showing the number of BGCs/gene clusters (X-axes) per GCF (Y-axes); (**2**) area plots showing the reduction in data set size (% of initial data set size, Y-axes) at each step in IGUA’s pipeline (X-axes). BGCs/gene clusters (GCs) correspond to the original input data set (i.e., 100% of the initial data set size), while FRs (fragment representatives), NRs (nucleotide representatives), and GCFRs (GCF representatives) correspond to the percentage of the input data set maintained at steps i (fragment removal), ii (nucleotide-level deduplication), and iii (protein-level clustering) in the IGUA pipeline, respectively; (**3**) BGC/gene cluster UMAPs of IGUA GCF representatives (points), colored by Seurat cluster; (**4**) histograms showing the number of phyla (X-axes) spanned by each GCF (Y-axes). For panels labeled 1 and 2, singletons have been excluded. For panels labeled 1 and 4, the X-axis maximum was set to 15.

### IGUA can be used for GCF delineation tasks beyond BGCs

IGUA’s iterative clustering approach is agnostic to protein function; unlike other BGC clustering tools, it does not rely on the presence of specific protein domains for GCF delineation. As such, IGUA can be used for GCF delineation tasks beyond BGC clustering. To demonstrate this, we used IGUA to delineate GCFs among two additional types of prokaryotic gene clusters: (i) secretion systems (*n* = 10,576 single-locus secretion systems from TXSSdb) (10); (ii) prophage (*n* = 356,776 prophage sequences from Prophage-DB) (51).

For (i) secretion systems, IGUA clustered all 10,576 single-locus secretion systems into 2,744 GCFs in under 3 minutes using 28 CPUs (Figure 3B, Supplementary Figure S6, Supplementary Tables S13-S15). Of all 2,744 GCFs, 2,639 (96.2%) contained a single secretion system type (singleton GCFs included; Supplementary Figure S6). In terms of their taxonomic spread, secretion system GCFs tended to be taxon-specific, as the vast majority were confined to a single phylum (2,421 of 2,744 GCFs, 88.2%, including singletons; Figure 3B, Supplementary Tables S13-S15). Within the secretion system UMAP, a total of 17 Seurat clusters could be discerned (clustering resolution = 0.35; Figure 3B). While some secretion system types were distributed across the UMAP (e.g., T5SS, T1SS to a lesser extent), others tended to be confined to a particular Seurat cluster (e.g., T2SS, T6SS; Figure 3B, Supplementary Figure S6).

Notably, T3SS clustered together with flagellar gene clusters (Figure 3B, Supplementary Figure S6), which is consistent with previous reports of a shared T3SS/flagellar genomic architecture and evolutionary history (56–58). Upon increasing the clustering resolution to the default 0.80, T3SS and flagellar gene clusters were split into two distinct clusters (Supplementary Figure S7). The UMAP further indicated that “tight adherence” (Tad) secretion systems formed a separate cluster, which was distinct from other T4SS (Figure 3B, Supplementary Figure S6). It has been hypothesized that bacterial Tad secretion systems were acquired from archaea via horizontal gene transfer (59–61), and our results are consistent with previous literature suggesting that Tad secretion systems are distinct from other bacterial secretion systems, including T4SS and type IV pili (T4P) (61).

Compared to the proGenomes3 BGC and TXSSdb single-locus secretion system datasets, which reduced to 2.0% and 25.9% of their original sizes, respectively (i.e., 2,829,071 BGCs to 56,960 GCFs and 10,576 secretion systems to 2,744 GCFs, respectively), (ii) prophage sequences were considerably more diverse: using a default clustering distance of 0.80, IGUA identified 213,699 GCFs among 356,776 prophage sequences, reducing Prophage-DB to 59.9% of its original size (Supplementary Table S16). Using an increased clustering distance threshold of 0.99 (out of a maximum of 1.0), IGUA produced 106,749 prophage GCFs (29.9% of the size of the original Prophage-DB; Figure 3C, Supplementary Figure S8, Supplementary Tables S17-S18). Like secretion systems, prophage GCFs tended to be confined to a single phylum (89,740 of 106,749 GCFs, 84.1%, including singletons), although a greater proportion of non-singleton prophage GCFs were distributed across multiple phyla (Figure 3C).

Overall, the GCF concept can be applied to biological problems beyond the BGC space. By generalizing GCF delineation, IGUA can provide novel insights into gene cluster ecology and evolution.

## DISCUSSION

Improvements in data generation methods, computing hardware, BGC mining software, and data storage/management practices are allowing BGC mining efforts to target larger (meta)genomic datasets than ever before (25, 39, 62). Scalable GCF delineation methods are thus crucial for making sense of these increasingly massive BGC datasets (27, 31). Here, we demonstrated that IGUA, our iterative GCF delineation method, has several advantages over previous methods designed specifically for BGC clustering. Most notably, we showed that IGUA is ≥10x faster than BiG-SLiCE, with comparable accuracy to BiG-SCAPE (the state-of-the-art methods in terms of speed and accuracy, respectively) (27, 31). Further, using 72 CPUs, we used IGUA to delineate GCFs among >2.8 million BGCs from nearly 1 million prokaryotic genomes in under 18 hours; for comparison, in the original publication (31), BiG-SLiCE took nearly 10 days to cluster >1.2 million BGCs (<50% of the BGCs queried in this study) into GCFs on a 36-core CPU server (50% of the CPUs used here). Finally, unlike BiG-SCAPE/BiG-SLiCE, which are specifically designed to cluster antiSMASH/MIBiG BGCs (28, 35), we show that IGUA can accept the output of alternative BGC detection tools (e.g., ML-based methods, such as GECCO) (21), as well as other types of gene clusters (here, single-locus secretion systems and prophage) (10, 51).

As demonstrated here, IGUA is particularly well-suited for large-scale BGC mining efforts (and large-scale gene cluster mining in general). However, it is important to point out the limitations of IGUA and outline scenarios/applications where users may prefer alternatives. First, unlike BiG-SCAPE (27), which includes a command-line option to cluster MIBiG BGCs alongside user-supplied BGCs, IGUA only clusters user-provided BGCs (or other gene clusters); thus, users must manually include MIBiG BGCs in their input GenBank file(s) if they wish for MIBiG to be included in an analysis. Second, BiG-SCAPE and BiG-SLiCE both produce graphical output files (e.g., interactive HTMLs summarizing a BiG-SCAPE/BiG-SLiCE run), while IGUA does not. Users who prefer graphical output formats may thus wish to use BiG-SCAPE/BiG-SLiCE, as no additional steps or analyses are needed to plot results. Finally, BiG-SCAPE and BiG-SLiCE are both embedded in the antiSMASH ecosystem, making them accessible and convenient for antiSMASH users (including those with limited bioinformatics experience). For example, a results directory produced by antiSMASH can be directly supplied as input to BiG-SCAPE (27), with minimal user intervention required. Similarly, many antiSMASH features (e.g., predicted biosynthetic classes of BGCs, comparison of antiSMASH BGCs to MIBiG BGCs) can be leveraged, reported, and/or summarized directly by BiG-SCAPE/BiG-SLiCE (IGUA, comparatively, only uses nucleotide/protein sequences for GCF delineation and thus does not consider any other information). Thus, antiSMASH users may appreciate the tailored antiSMASH integration that BiG-SCAPE and BiG-SLiCE provide.

We anticipate that IGUA will be particularly useful for e.g., redundancy removal (particularly among gene clusters detected in highly fragmented MAGs), functional prediction of BGCs, and novel BGC discovery/prioritization, particularly among users of increasingly popular ML-based BGC discovery tools (21, 22, 24, 25). We further posit that large-scale (meta)genomic mining efforts targeting other types of prokaryotic gene clusters (e.g., secretion systems, prophage, genomic islands, phage defense systems) (10, 18, 51) will benefit from IGUA’s flexibility, as IGUA can be used for any type of genomic sequence segment clustering problem. Altogether, IGUA represents a versatile GCF delineation tool with unmatched computational efficiency and flexibility, enabling (meta)genomic mining applications at unprecedented scales.

## DATA AVAILABILITY

IGUA is open-source and freely available via GitHub under the GPL-3.0 License (https://github.com/zellerlab/IGUA). Source code for the IGUA version used in the manuscript (v0.1.0; i.e., the IGUA version used in the manuscript, but with default parameters set to the optimal values identified in this study) is available via Zenodo (https://doi.org/10.5281/zenodo.15348788). Source and compiled releases are available to download from the Python Package Index (PyPI, https://pypi.org/project/igua/) and from Bioconda (https://anaconda.org/bioconda/igua) (63). Accession numbers of genomes used in the study, as well as all benchmarking and validation results generated in the study, are available as Supplementary Data. Analysis code is available via GitHub (https://github.com/c20josbl/igua_manuscript_documentation).

## SUPPLEMENTARY DATA

Supplementary Data are available via Zenodo under DOI: 10.5281/zenodo.15394038 (https://doi.org/10.5281/zenodo.15394038).

## AUTHOR CONTRIBUTIONS

Software development was performed by ML, with input from LMC. Computational analyses were performed by JB, with contributions from ML, HG, and LMC. ML, LMC, and GZ conceived the study, which was funded by LMC and GZ. JB, ML, and LMC co-wrote the manuscript with contributions from all authors.

## ACKNOWLEDGMENTS

This research was conducted using the resources of High Performance Computing Center North (HPC2N; Umeå University, Umeå, Sweden), as well as the HPC resources of the European Molecular Biology Laboratory (EMBL, Heidelberg, Germany).

