## Supplementary Figure S1 for "Fast, flexible gene cluster family delineation with IGUA"

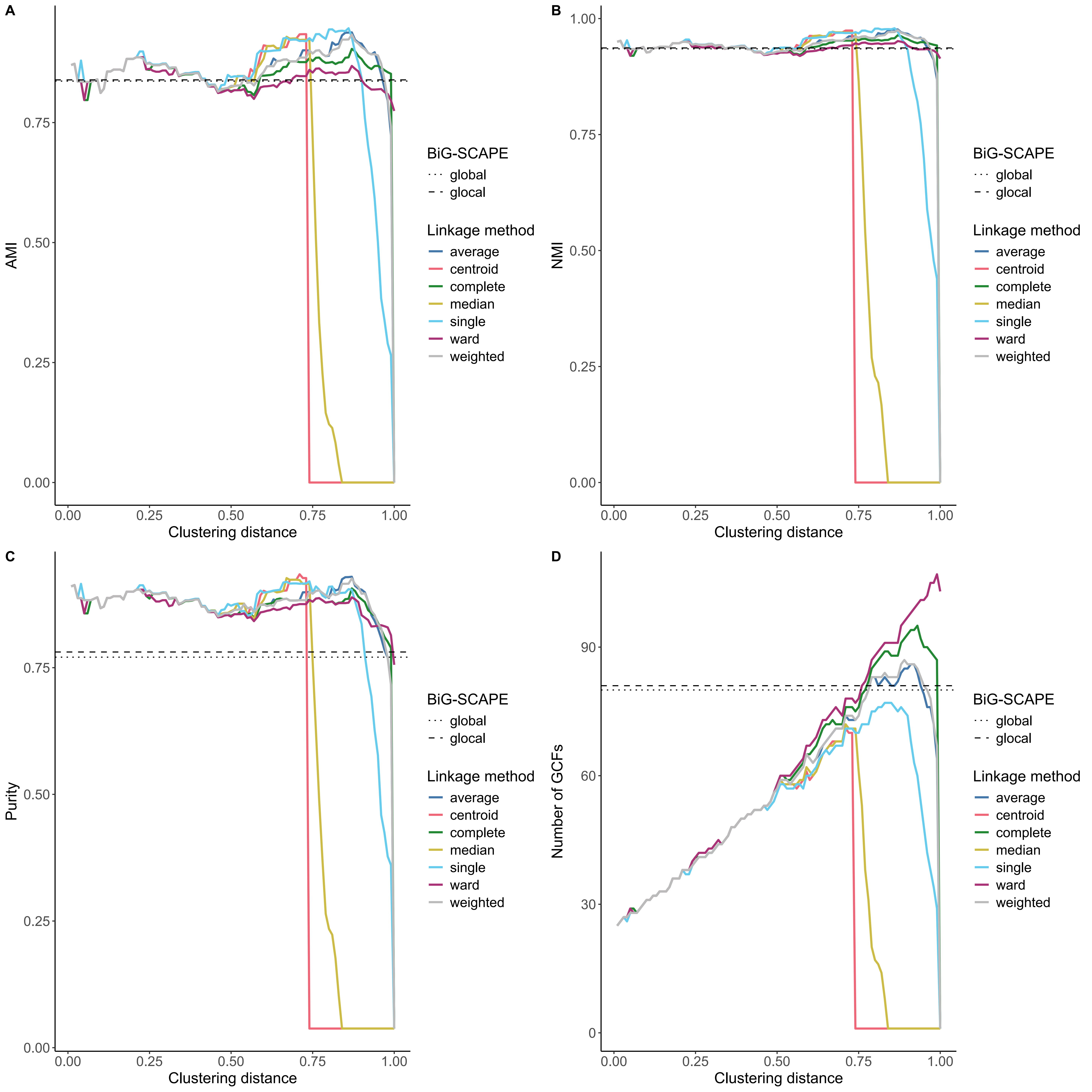


Supplementary Figure S1. (A) Adjusted mutual information (AMI), (B) normalized mutual information (NMI), (C) purity, and (D) number of Gene Cluster Families (GCFs; Y-axes) of IGUA GCFs relative to manually curated, “ground truth” GCFs for each tested parameter combination (clustering distance, X-axis; hierarchical clustering linkage method, line color). Horizontal dashed lines denote values achieved by BiG-SCAPE in “global” or “glocal” mode. A clustering distance of 0.80 and the “average” linkage method were selected as IGUA’s default parameters.
