## Supplementary Figure S2 for "Fast, flexible gene cluster family delineation with IGUA"

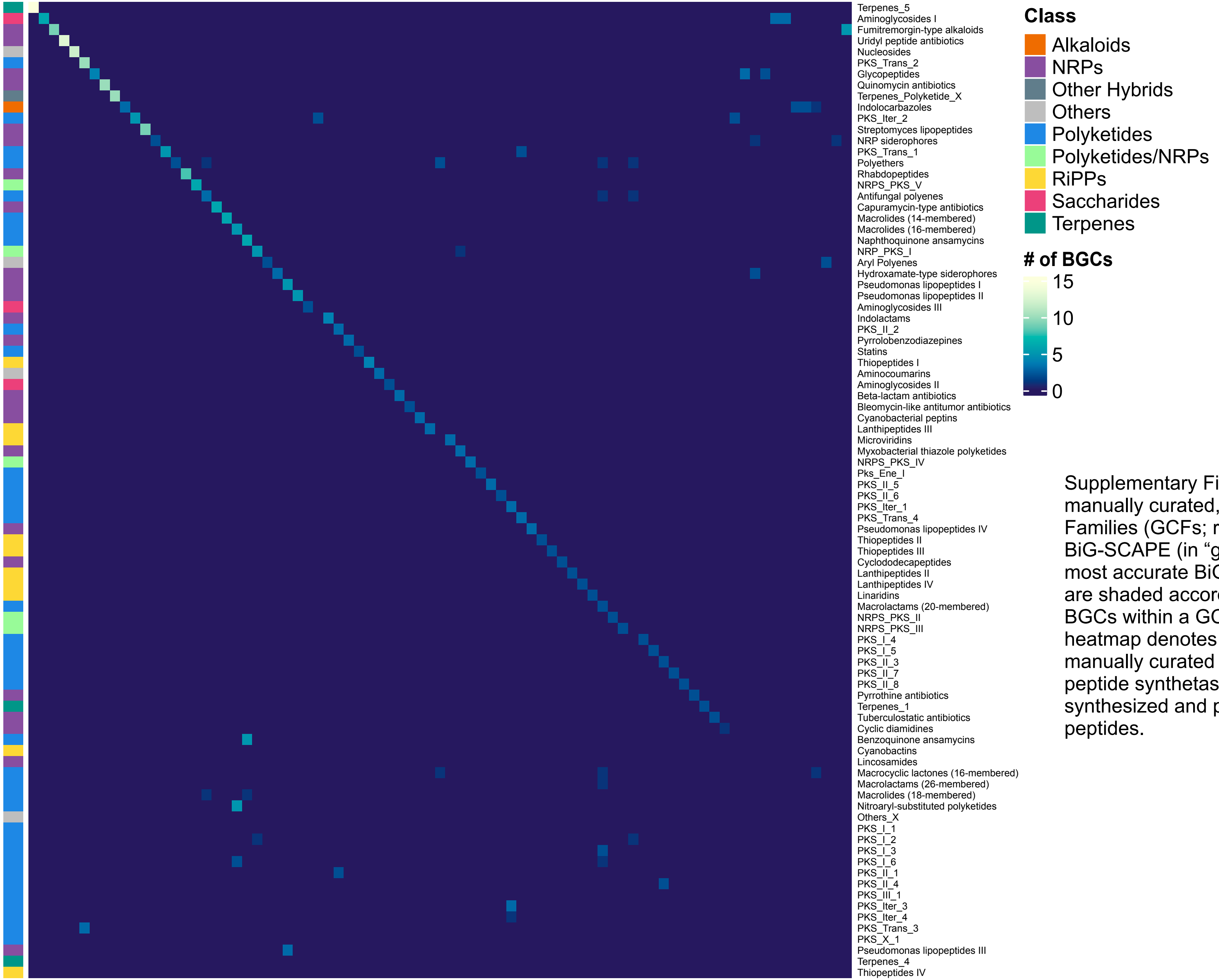


Supplementary Figure S2. Heatmap comparing manually curated, "ground truth" Gene Cluster Families (GCFs; rows) to GCFs delineated using BiG-SCAPE (in "glocal" mode, which produced the most accurate BiG-SCAPE results; columns). Cells are shaded according to the number of overlapping BGCs within a GCF. The color strip to the left of the heatmap denotes the biosynthetic class of each manually curated GCF. NRPs, non-ribosomal peptide synthetases; RiPPs, ribosomally synthesized and post-translationally modified peptides.
