## Supplementary Figure S3 for "Fast, flexible gene cluster family delineation with IGUA"

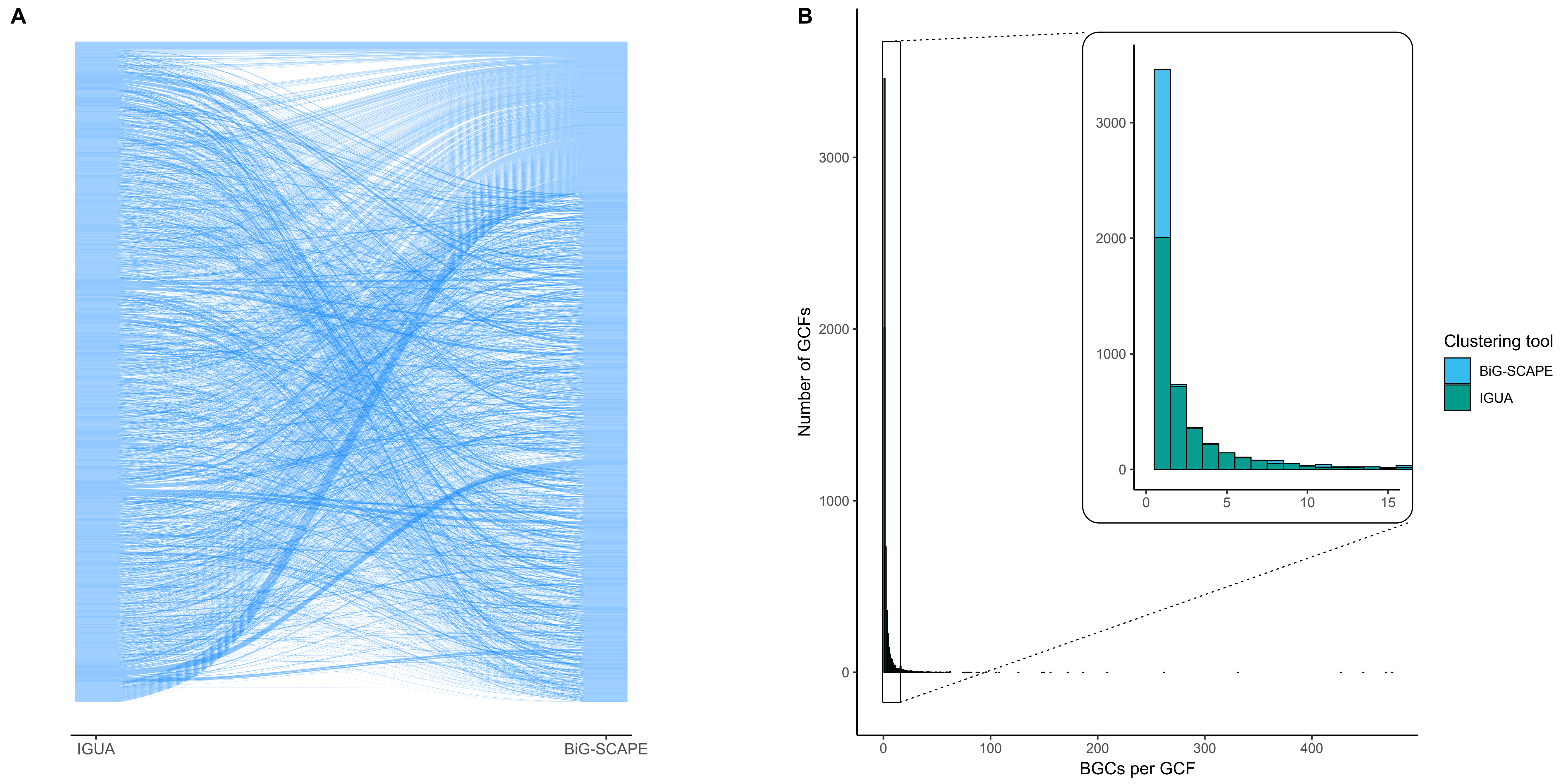


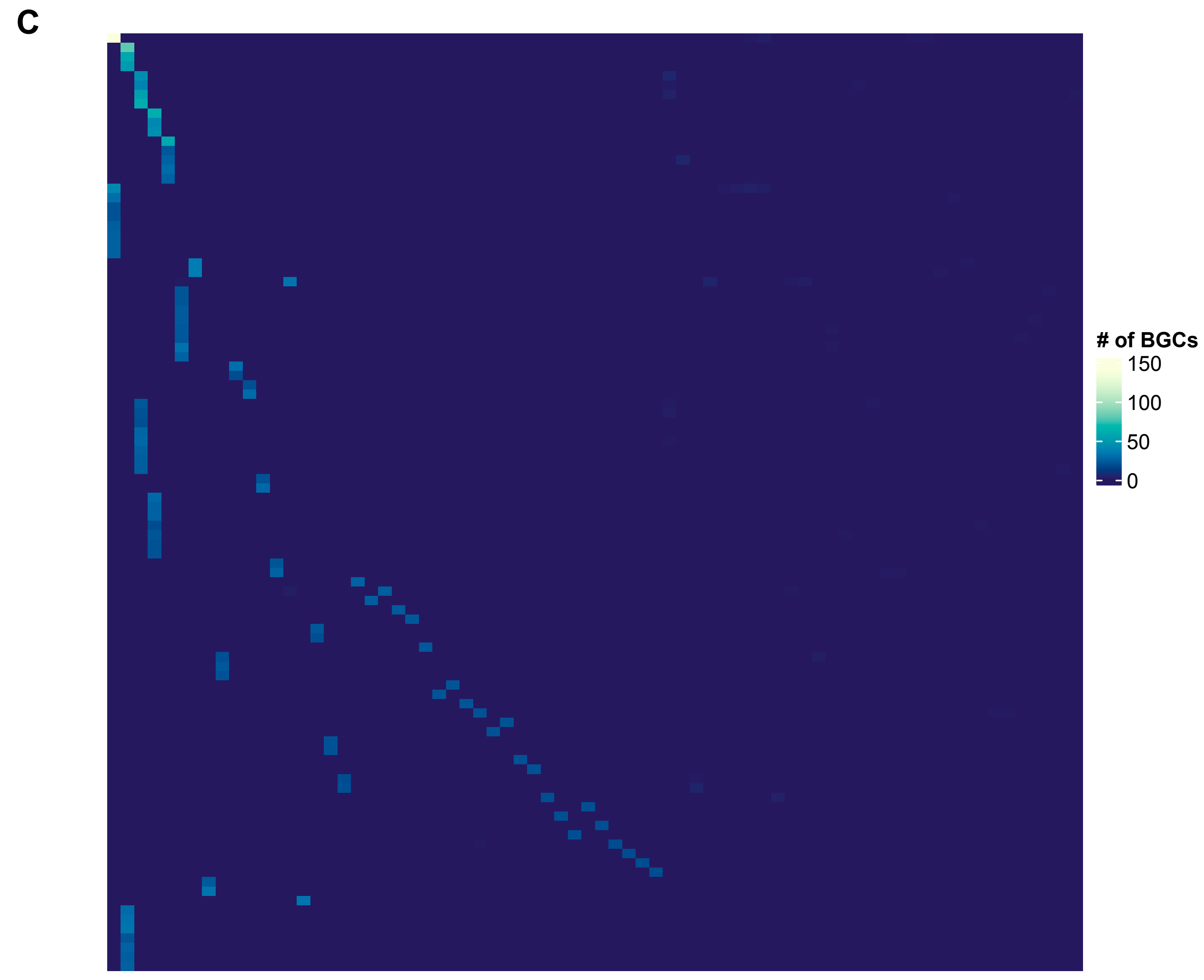


| **Metrics** | With singletons | Without singletons |
|---|---|---|
| **AMI** | 0.533 | 0.612 |
| **ARI** | 0.826 | 0.823 |
| **NMI** | 0.199 | 0.211 |

Supplementary Figure S3. Comparison of IGUA and BiG-SCAPE GCFs on a Streptomyces dataset. (A) Alluvial plot comparing the composition of Streptomyces GCFs delineated by IGUA (left) and BiG-SCAPE (right). The plot was constructed using the geom_alluvium function from the ggalluvial package. (B) Histogram of the number of BGCs per Streptomyces GCF (X-axis) detected using BiG-SCAPE (blue) and IGUA (green). (C) Heatmap comparing GCFs delineated using BiG-SCAPE (rows) and IGUA (columns). For readability, only the 100 largest overlapping GCFs are shown. Metrics to the right of the heatmap pertain to the entire Streptomyces dataset. AMI, adjusted mutual information; ARI, adjusted Rand index; NMI, normalized mutual information.
