## Supplementary Figure S4 for "Fast, flexible gene cluster family delineation with IGUA"

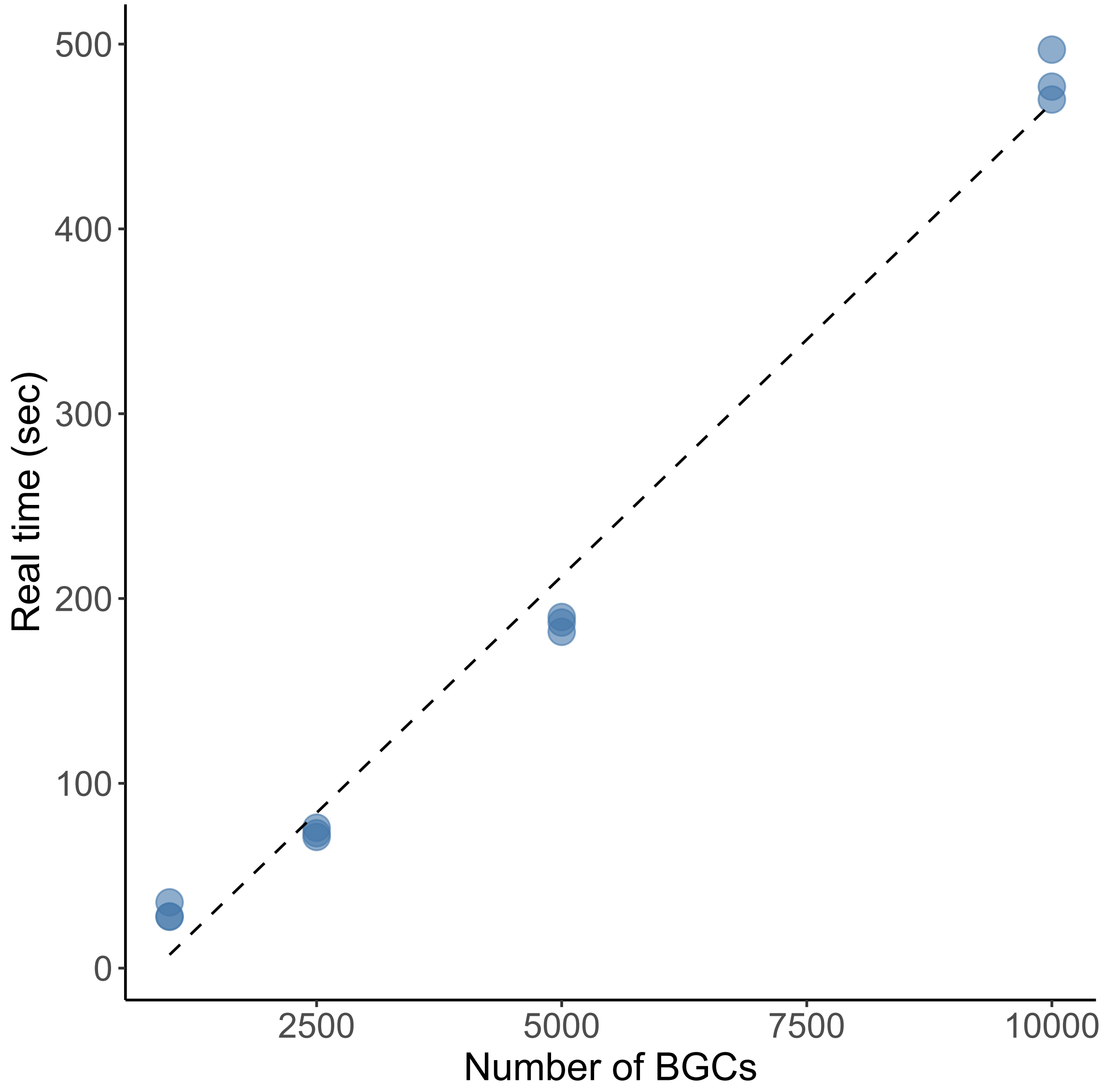


Supplementary Figure S4. Real/“wall clock” time (in seconds; Y-axis) needed to cluster (i) 1,000, (ii) 2,500, (iii) 5,000 and (iv) 10,000 BGCs (X-axis) using IGUA (default settings, 8 CPUs). BGCs were randomly sampled in triplicate from a set of Streptomyces antiSMASH BGCs, with MIBiG BGCs included. Points denote IGUA’s runtime for each randomly sampled BGC set, while the dashed line denotes the best-fitting linear model, constructed using the following: geom_smooth(method = "glm", colour="black", lwd=0.5, linetype=2, se=FALSE).
