## Supplementary Figure S5 for "Fast, flexible gene cluster family delineation with IGUA"

A

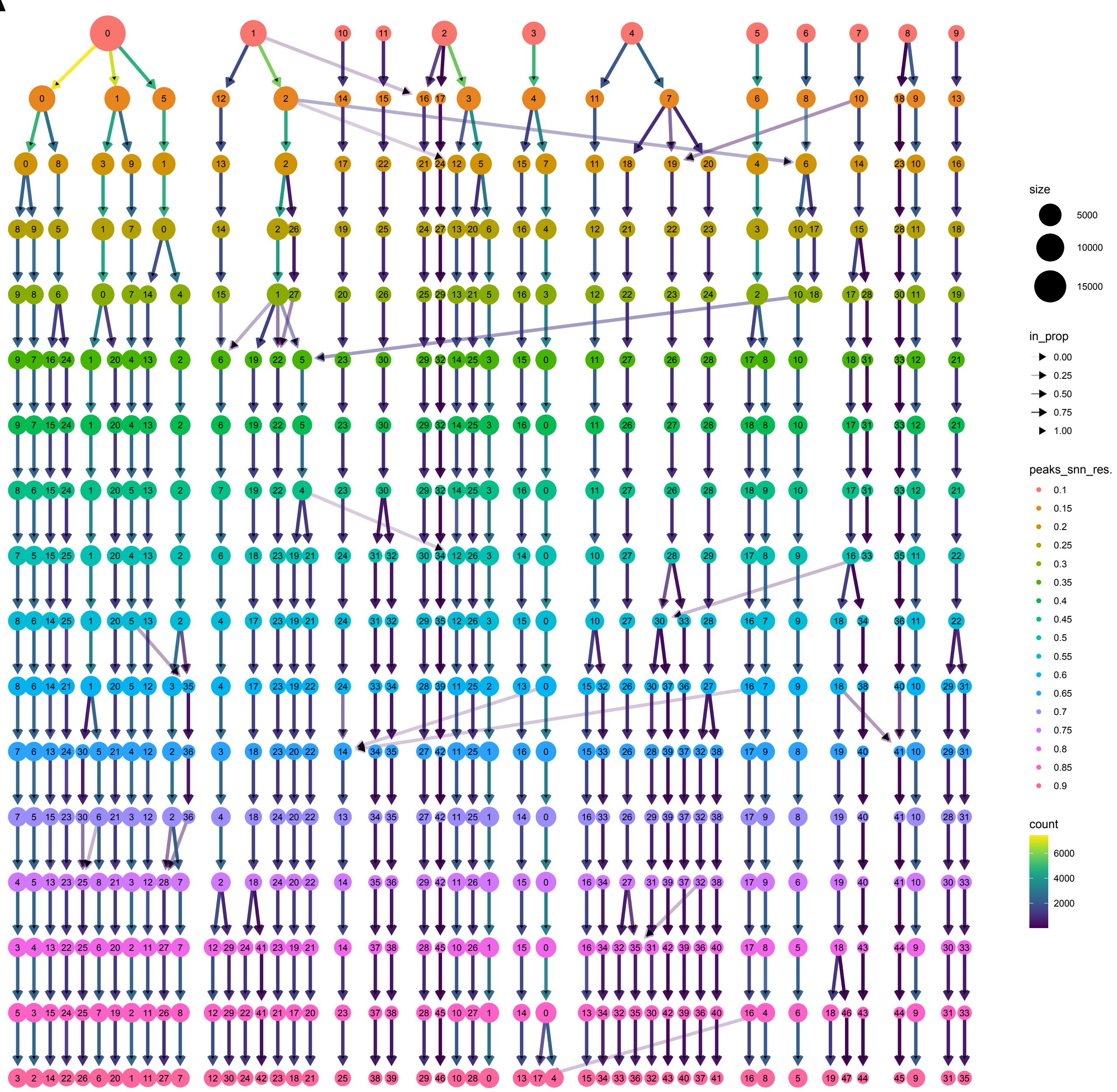


Supplementary Figure S5. (A) Clustering tree for the proGenomes3 UMAP. (B) proGenomes3 UMAP, colored by predicted biosynthetic class of the majority of BGCs in the GCF. Each dot represents a Gene Cluster Family (GCF) representative identified using IGUA.

B

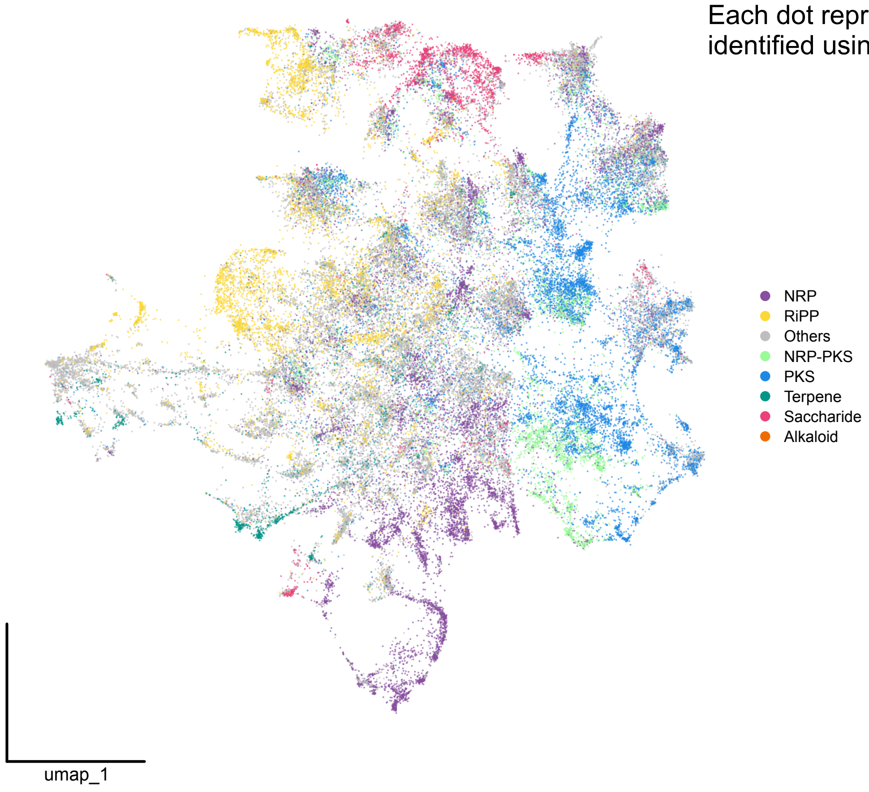
