## Supplementary Figure S6 for "Fast, flexible gene cluster family delineation with IGUA"

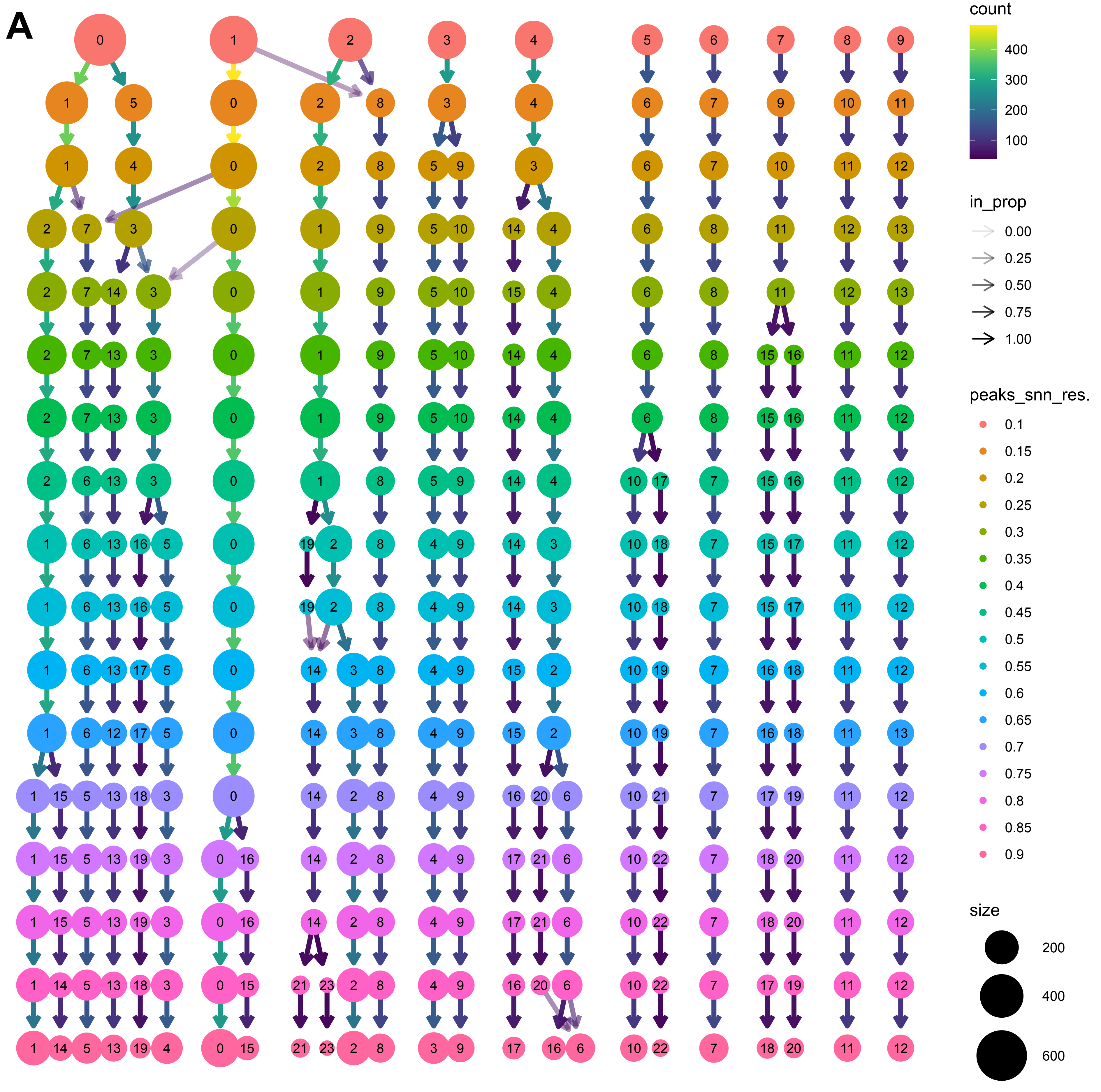


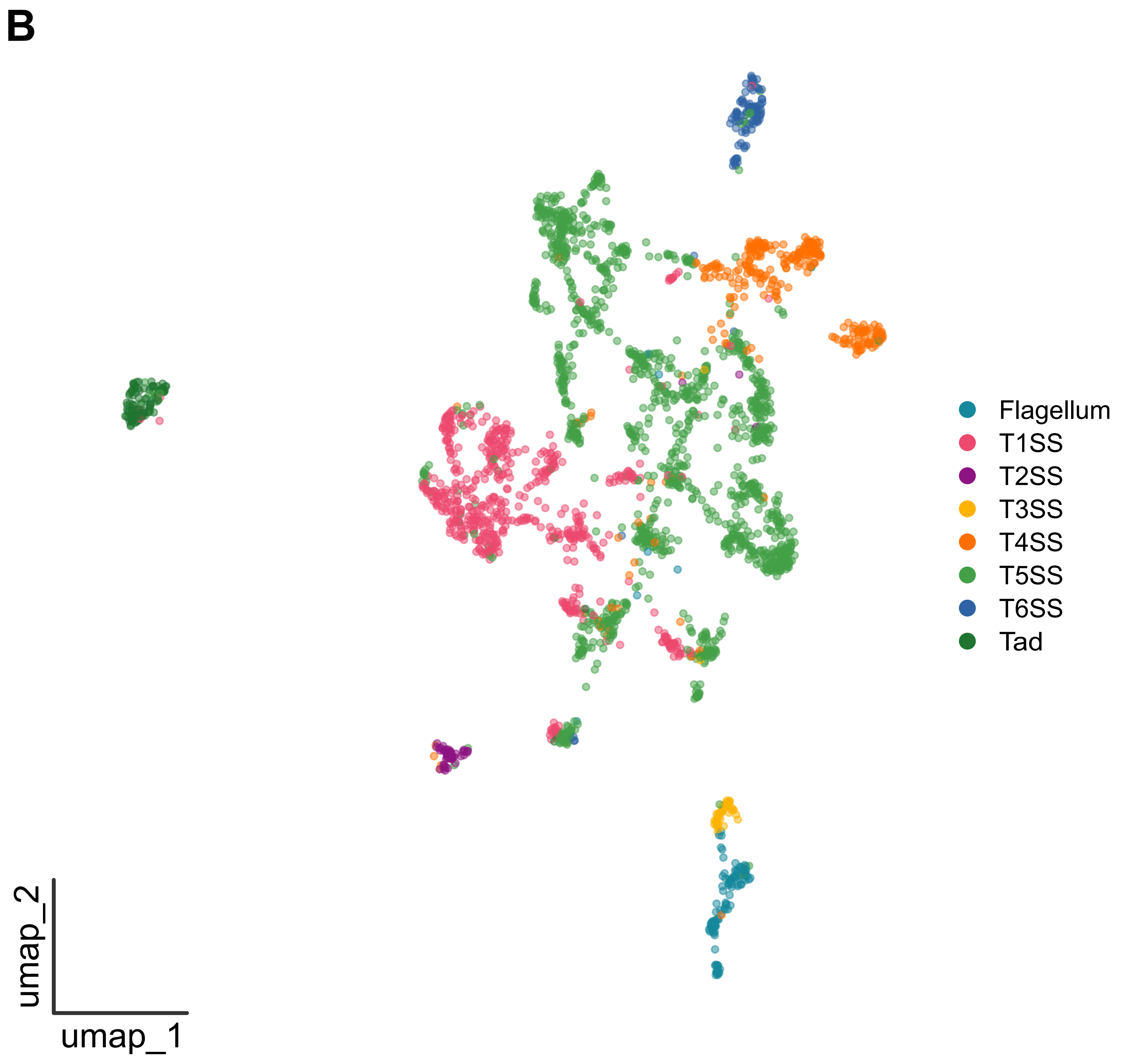


Supplementary Figure S6. (A) Clustering tree for (B) the TXSSdb single-locus secretion system UMAP. For (B), the secretion system UMAP is colored by secretion system type (provided by TXSSdb).
