## Supplementary Figure S7 for "Fast, flexible gene cluster family delineation with IGUA"

Supplementary Figure S7. The TXSSdb single-locus secretion system UMAP, colored by Seurat clusters obtained at a default clustering resolution of 0.80. The red circle denotes Seurat clusters containing T3SS and flagellar gene clusters.

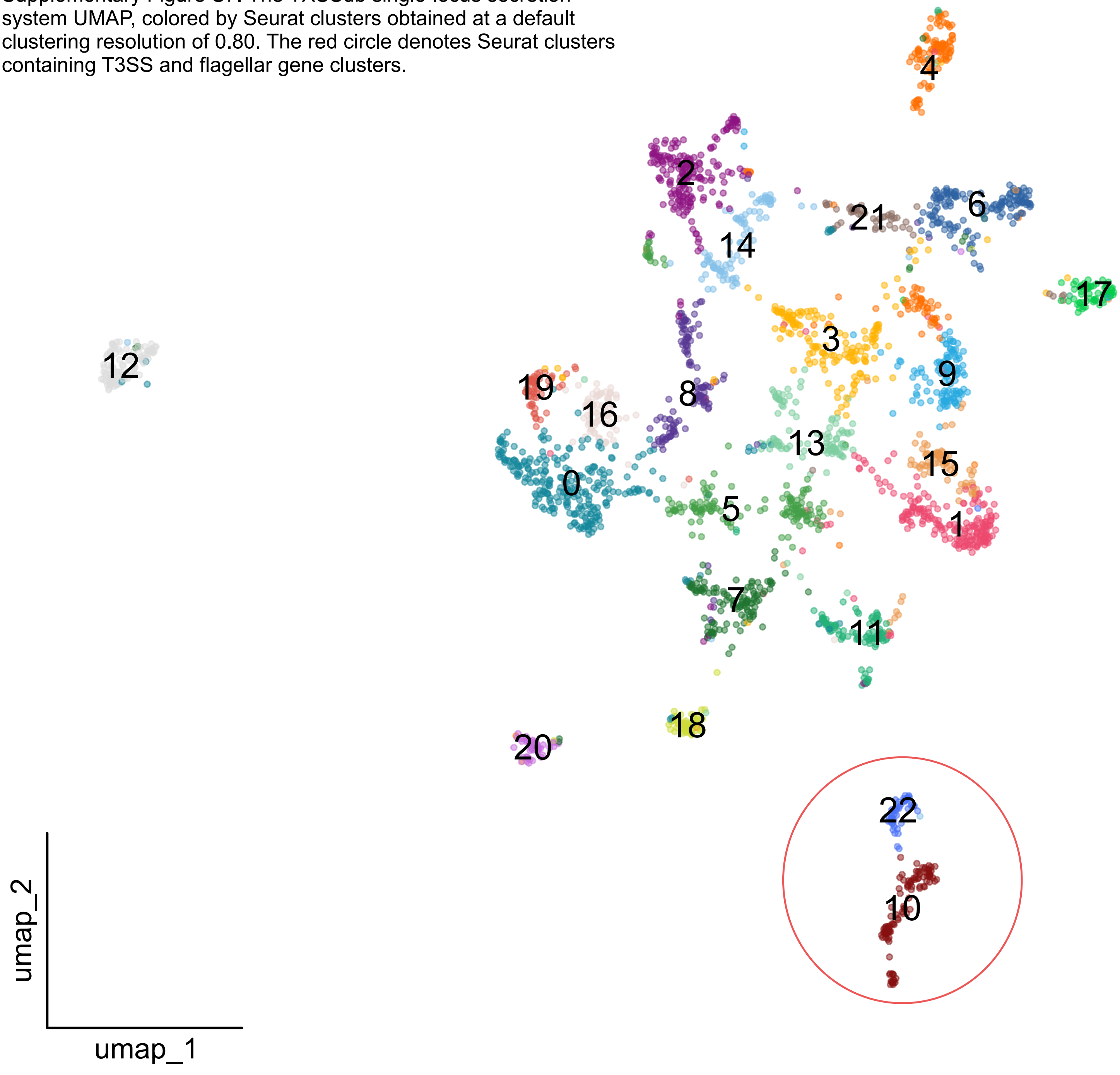
