## Supplementary Figure S8 for "Fast, flexible gene cluster family delineation with IGUA"

A

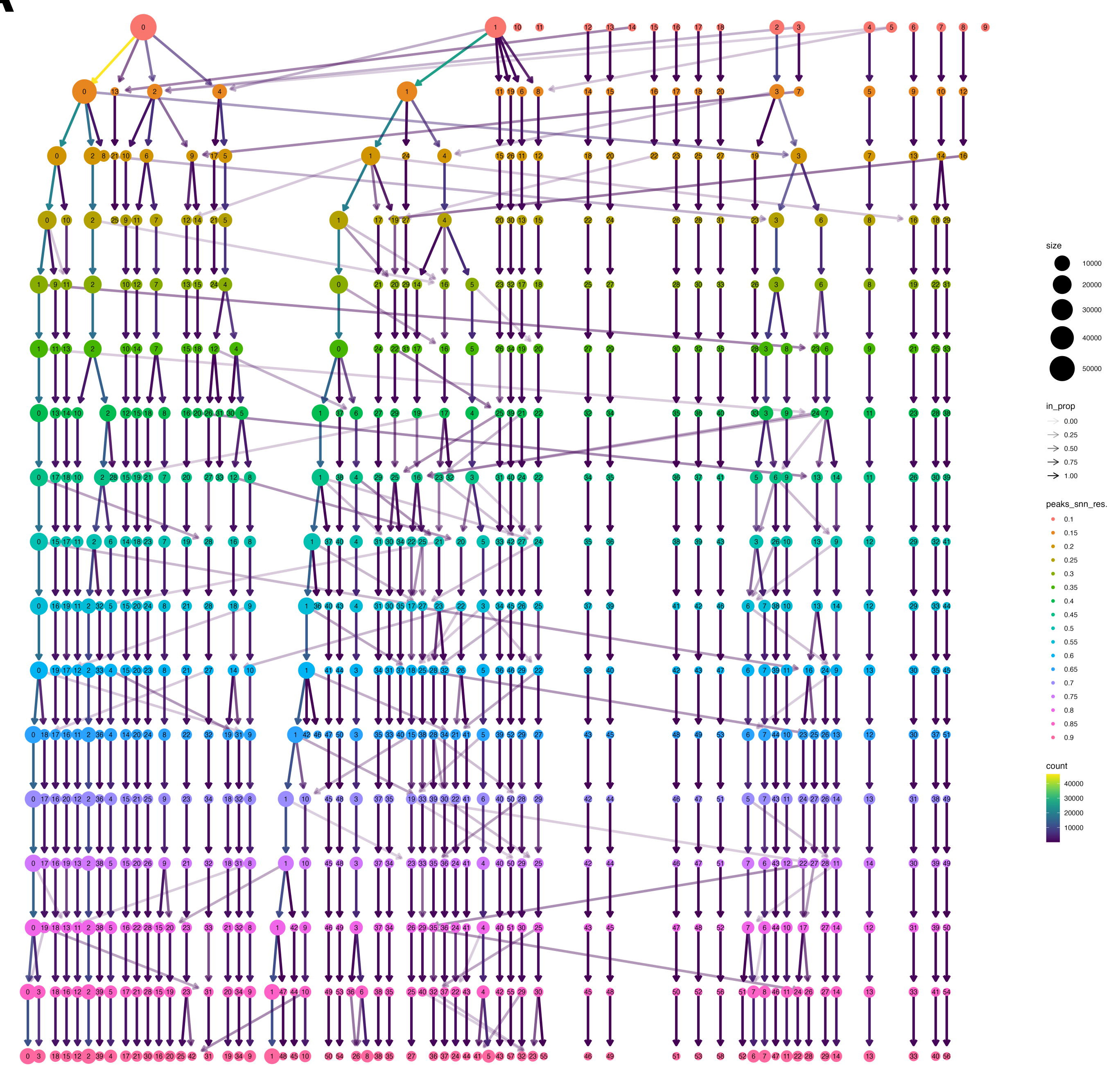


B

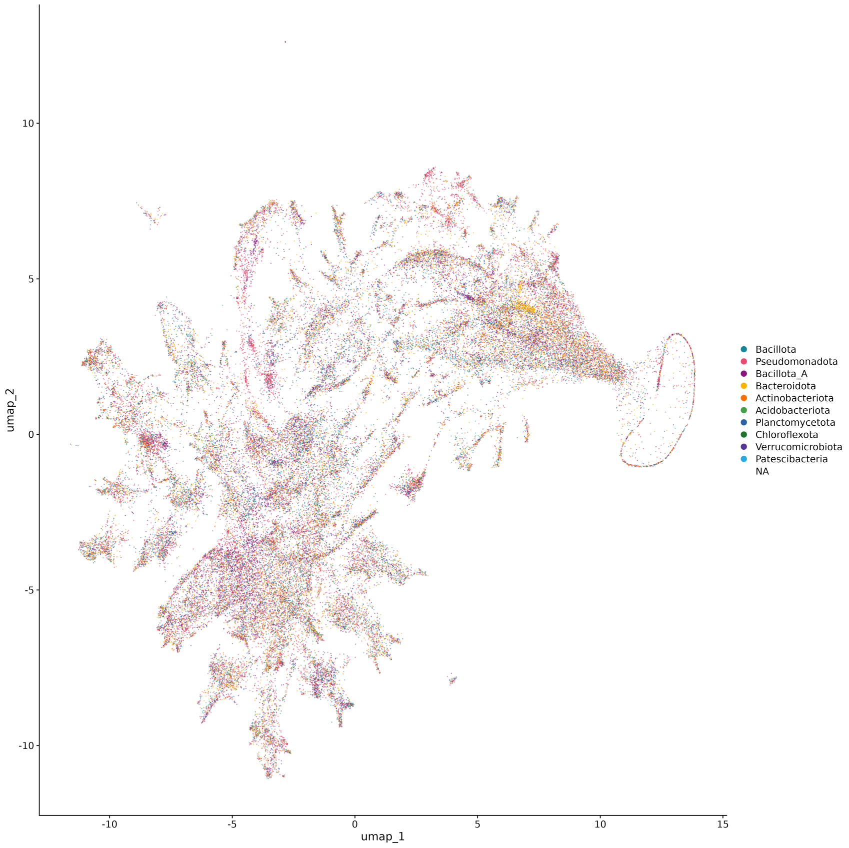


Supplementary Figure S8. (A) Clustering tree for (B) the Prophage-DB prophage UMAP, colored by phylum. For (B), only the top 10 phyla are shown (the rest are labeled as “NA”).
